# A ubiquitination-mediated degradation system to target phospho-14-3-3-binding-motif embedded proteins

**DOI:** 10.1101/2022.11.22.517526

**Authors:** Zhaokai Li, Xiaoqiang Huang, Mohan Li, Ziad Sabry, Y. Eugene Chen, Zhong Wang, Liu Liu

## Abstract

The phosphorylation of 14-3-3 binding motif is involved in many cellular processes. A strategy that enables targeted degradation of phospho-14-3-3-binding-motif embedded (P-14-3-3-BME) proteins for studying their functions is highly desirable for basic research. Here, we report a phosphorylation-induced, ubiquitin-proteasome-system-mediated targeted protein degradation (TPD) strategy that allows specific degradation of P-14-3-3-BME proteins. Specifically, by ligating a modified von Hippel-Lindau E3-ligase with an engineered 14-3-3 bait, we generated a protein chimera referred to as Targeted Degradation of P-14-3-3-BME Protein (TDPP). TDPP can serve as a universal degrader for P-14-3-3-BME proteins based on the specific recognition of the phosphorylation in 14-3-3 binding motifs. TDPP shows high efficiency and specificity to a difopein-EGFP reporter, general and specific P-14-3-3-BME proteins. TDPP can also be applied for the validation of P-14-3-3-BME proteins. These results strongly support TDPP as a powerful tool for 14-3-3 related research.

## Introduction

Controlling a protein’s concentration level is key to studying its function. Strategies that specifically target and degrade a protein of interest (POI) have emerged in recent years, which are collectively called targeted protein degradation (TPD) (Ding et al., 2020). Among them, strategies based on the ubiquitin-proteasome system (UPS), a crucial protein degradation system in eukaryotic cells (Pickart, 2001; Zheng and Shabek, 2017), including heterobifunctional small molecules like proteolysis targeting chimeras (PROTACs) (Schneekloth et al., 2008), protein/peptide-based degraders like affinity-directed protein missiles (AdPROMs) (Fulcher et al., 2016), molecular glue (Tan et al., 2007), dTAG (Nabet et al., 2018), etc., show great efficiency for protein degradation..

However, current TPD strategies still have their limitations. Only a few tag-based TPD strategies (Buckley et al., 2015; Nabet *et al*., 2018; Neklesa et al., 2011) (reviewed by Burslem, et al. (Burslem and Crews, 2020)) have been developed to target proteins with post-translational modifications (PTMs), while specific TPD strategies are still lacking for major types of PTMs that widely exist in eukaryotic cells, including protein phosphorylation, methylation, acetylation, etc., which play an important role in regulating protein functions. The same protein under different PTM statuses can display different functions. Therefore, the development of PTM-specific degradation systems is desirable.

The TPD strategy has the potential to regulate specific kinds of phosphorylated proteins. While at least three quarters of the proteome is phosphorylated (Sharma et al., 2014), protein phosphorylation exists in every organism and participates in most biological events (Ardito et al., 2017). The phosphorylation/dephosphorylation states of proteins can be distinguished by their “readers” that interact with the binding motifs. 14-3-3 is one of those phosphorylation readers. As one of the most expressed proteins in eukaryotic cells (Wang et al., 2015), 14-3-3 is a highly conserved protein family consisting of 7 isoforms (i.e., β, γ, ε, ζ, η, τ, and σ) (Cornell and Toyo-Oka, 2017). The function of 14-3-3 largely depends on its interaction with phospho-serine/threonine (phospho-S/T) in phosphoproteins (Fu et al., 2000), which are embedded within phospho-14-3-3 binding motifs.

Here we describe a new TPD system, namely TDPP, for Targeted Degradation of Phospho-14-3-3-binding-motif-embedded (P-14-3-3-BME) Protein. TDPP is a protein chimera made by coupling a modified von Hippel-Lindau (VHL) E3-ligase with an engineered 14-3-3ζ bait. Independent experiments showed that TDPP targets P-14-3-3-BME proteins with high affinity and efficiency. Furthermore, with high specificity to target the proteins of interest, it will likely be a powerful biological tool for protein-protein interaction (PPI) research, regulation of the phosphorylation/dephosphorylation status of multiple proteins and the level of phospho-proteins at the whole-cell level.

## Results

### Design of a degradation system targeting P-14-3-3-BME proteins

Inspired by previously described hetero bifunctional degrader strategies (Burslem and Crews, 2020; Caussinus et al., 2011; Crew et al., 2018; Fulcher et al., 2017; Ibrahim et al., 2020), we developed a degradation system that specifically targets P-14-3-3-BME proteins (Fig. 1A). Following the logic that a 14-3-3 bait is appropriate to recruit the P-14-3-3-BME protein targets and an E3 ligase is necessary to introduce the POI to UPS for degradation, an E3-14-3-3 ligated protein became the fundamental structure of the degrader described here. While VHL is an E3 ligase that numerous TPD strategies were based on, and 14-3-3ζ ranks as one of the major 14-3-3 isoforms in humans, those two were picked as the UPS-binding part and the POI-binding part, respectively. It was reported that the C-terminal of wildtype (WT) 14-3-3 protein isoforms decrease their binding efficiency to 14-3-3 binding motif (Truong et al., 2002), so a C-terminal truncated version of 14-3-3ζ was used, i.e., 14-3-3ζ was truncated to 14-3-3ζ (1-230). Finally, giving to the protein nature of the protein chimera described above, which would potentially lead to auto-ubiquitination and auto-degradation when it is exposed to UPS, all the lysine (K) residues within the protein chimera were mutated to arginine (R) following a proven strategy (Yin et al., 2019), and VHL 3KR-14-3-3ζ (1-230) 20KR (referred to as TDPP, Fig. 1, B and C) was finally developed and evaluated here after.

**Figure 1.**
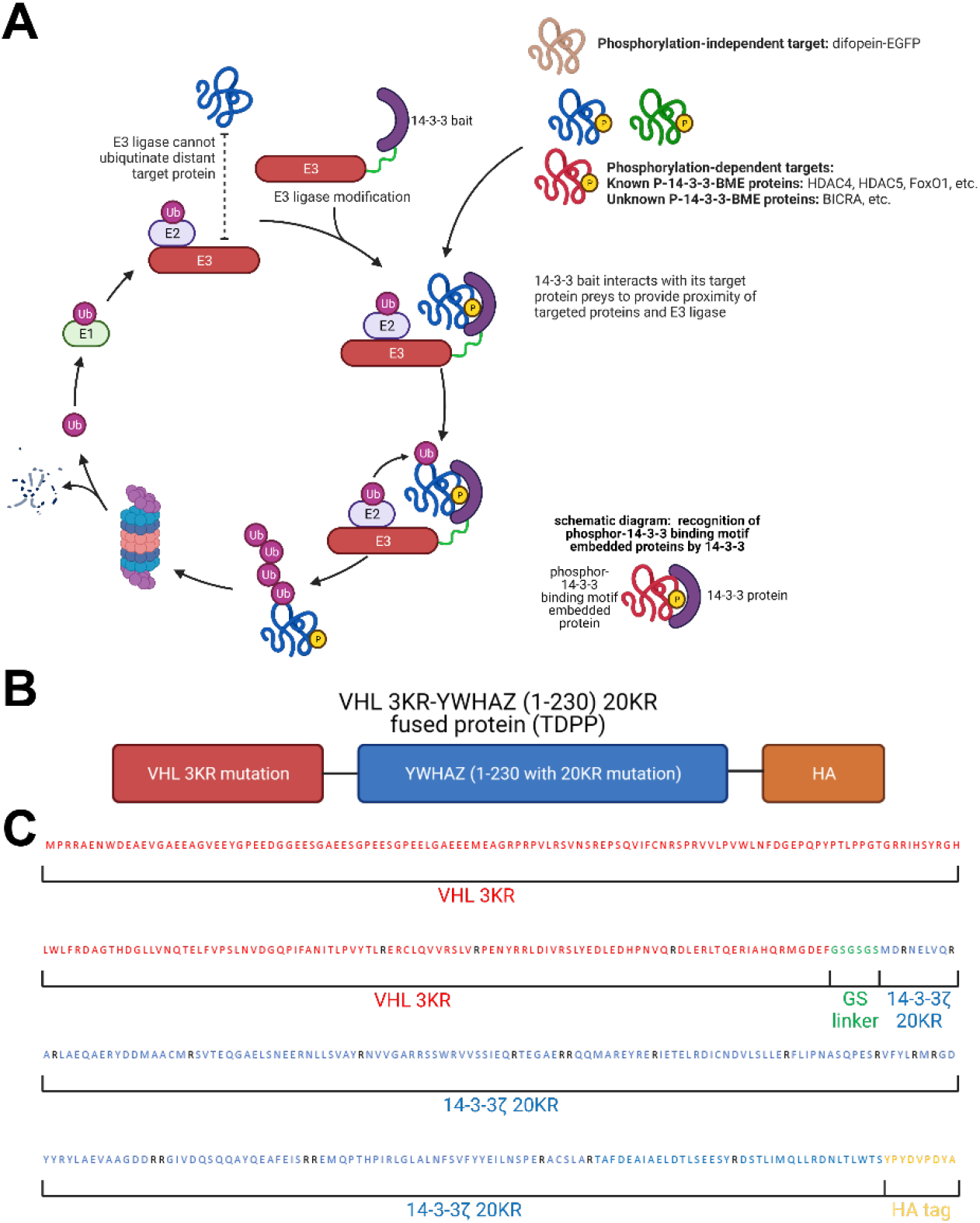
Graphic summary and illustration of a degrader of phospho-14-3-3-binding-motif embedded proteins. A. Working model of TDPP to degrade P-14-3-3-BME proteins. B. TDPP fused protein design. C. Sequence partition information of TDPP.

A protein’s structure determines its function. To know if the domains of the protein chimera [i.e., VHL 3KR and 14-3-3ζ (1-230) 20KR] in TDPP could properly fold into desired structures, we used ColabFold (Mirdita et al., 2022) which is adapted from AlphaFold2 (Jumper et al., 2021) to perform de novo structure prediction of TDPP without any structure templates. Specifically, we directly submitted the TDPP protein sequence to the Colab Fold server (https://colab.research.google.com/github/sokrypton/ColabFold/blob/main/AlphaFold2.ipynb) and chose to predict five models with AMBER refinement. All other parameters were used as default. The average predicted local-distance difference test (pLDDT) scores for all five models range from 81.2 to 82.7 (Supplementary Fig. 1B), indicating the overall high quality of the models [usually an LDDT score of 0.6 or greater is considered a reasonable model and scores above 0.8 are great models (Mariani et al., 2013)]. The relatively low per-residue LDDT scores suggest that the N-terminal of VHL, the GSGSGS linker, and the hemagglutinin (HA) tag regions are of lower quality (Supplementary Fig. 1A), consistent with the annotation in UniProt that the N-terminal (i.e., amino acids 1-65) of VHL is disordered. Otherwise, the other regions of VHL and 14-3-3ζ are well folded (Supplementary Fig. 1C). Except the second-ranking model (i.e., *rank_2*), the other four models have their N-terminal tail inserted into the binding groove of 14-3-3ζ (Supplementary Fig. 1C). However, no evidence supports the binding between the N-terminal of VHL and 14-3-3ζ and the insertions are possibly caused by the less accurate modeling of the VHL N-terminus. Fig. 2A depicts the overall structure of TDPP (*rank_2* is used). As shown in Fig. 2, B and C, the predicted VHL and 14-3-3 protein components match perfectly with their native counterparts respectively, except for the disordered region. Taken together, de novo structure modeling suggests that the individual components of TDPP can fold properly to perform desired function.

**Figure 2.**
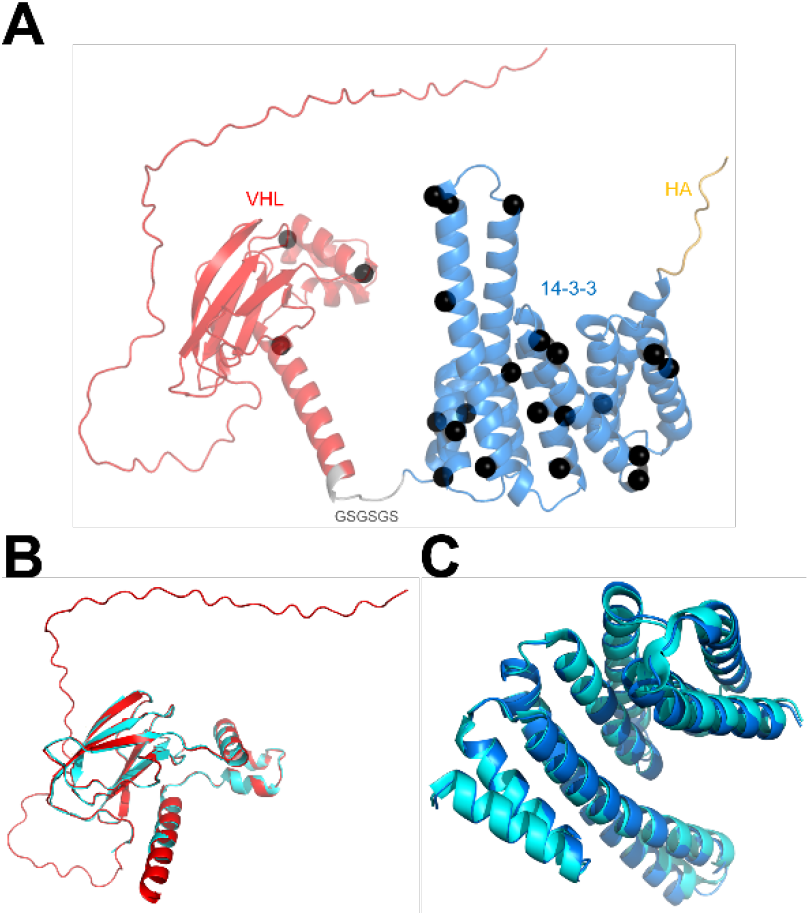
De novo structure modeling of TDPP. A. Full-length structure model generated by ColabFold(Mirdita *et al*., 2022). KR mutation sites are highlighted in black spheres. B. Comparison of predicted VHL 3KR model and experimental VHL structure. Red: predicted VHL (extracted from full-length model); cyan: experimental VHL structure (PDB ID: 4WQO). C. Comparison of predicted 14-3-3ζ 20 KR model and experimental 14-3-3ζ structure. Blue: predicted 14-3-3ζ (extracted from full-length model); cyan: experimental 14-3-3ζ structure (PDB ID: 2O02).

### TDPP degrades difopein-EGFP

The fundamental hypothesis of TDPP as a TPD strategy is that TDPP could interact with P-14-3-3-BME proteins when they are phosphorylated, and their interaction with TDPP could lead to the degradation of the interacting proteins themselves. Because of the technical difficulty to control the dynamic change of the phosphorylation/dephosphorylation status of P-14-3-3-BME proteins in cells, the process to evaluate TDPP’s efficiency was divided into two separate parts: whether TDPP could induce the degradation of its interacting partners and whether TDPP could induce the degradation of P-14-3-3-BME proteins. The former question was assessed with a phosphorylation-independent evaluation system. A small peptide inhibitor of 14-3-3 protein, difopein (Masters and Fu, 2001; Wang et al., 1999), was fused to an EGFP fluorescence reporter. Such a difopein-EGFP fused protein could become an excellent POI of TDPP strategy, which mimics the binding of 14-3-3-BME proteins to 14-3-3 and holds a reliable interaction ability with the TDPP chimera, all while the interaction is phosphorylation-independent (Masters and Fu, 2001). Thus, the 14-3-3-difopein interaction was expected to induce stable difopein-EGFP degradation and therefore lead to declining EGFP fluorescence (Fig.1A and Fig. 3A). Both the TDPP chimera and difopein-EGFP fused protein were expressed in HEK293T/17 cells using the co-transfection strategy (see Methods for details). As predicted, we observed a significantly decreased EGFP protein level when TDPP was introduced, compared with the control group (Fig. 3, B and C). In parallel, we also observed a significant decrease in EGFP fluorescence for the TDPP-treated group (Fig. 3, D and E).

**Figure 3.**
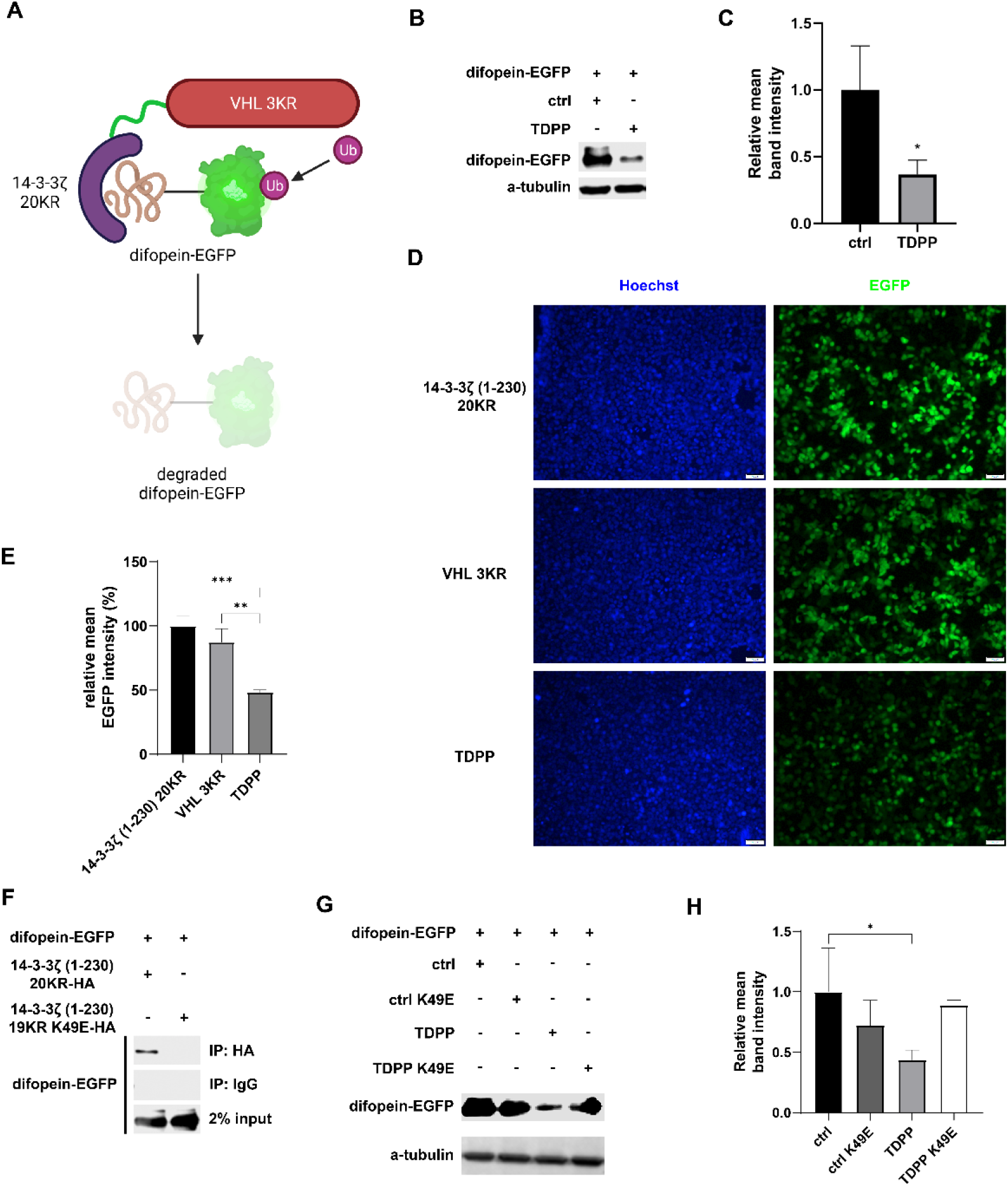
TDPP degrades difopein-EGFP. A. Working model of TDPP-induced degradation of difopein-EGFP. B, C. Western blot result (B) and quantification of band intensity (C) of difopein-EGFP in HEK293T/17 cells co-transfected with expression plasmids of 14-3-3ζ (1-230) 20KR-HA (referred as ctrl) or VHL 3KR-14-3-3ζ (1-230) 20KR-HA (referred as TDPP). D, E. Fluorescence assay result (D) and fluorescence intensity quantification (E) of HEK293T/17 cells transfected with difopein-EGFP vector, along with control vectors (14-3-3ζ (1-230) 20KR or VHL 3KR) or TDPP vector. F. Immunoprecipitation result of interaction between 14-3-3ζ (1-230) 20KR-HA/14-3-3ζ (1-230) 19KR K49E-HA and difopein-EGFP. IP: 14-3-3ζ (1-230) 20KR-HA/14-3-3ζ (1-230) 19KR K49E-HA. IB: difopein-EGFP. G, H. Western blot result (G) and quantification of band intensity (H) of difopein-EGFP in HEK293T/17 cells co-transfected with expression plasmids of 14-3-3ζ (1-230) 20KR-HA (referred as ctrl), 14-3-3ζ (1-230) 19KR K49E-HA (referred as ctrl K49E), VHL 3KR-14-3-3ζ (1-230) 20KR-HA (referred as TDPP), or VHL 3KR-14-3-3ζ (1-230) 19KR K49E-HA (referred as TDPP K49E).

We next determined whether the decreased difopein-EGFP phenotype was caused by its interaction with TDPP. Since lysine 49 (K49) residue of 14-3-3 plays a key role in interacting with difopein or P-14-3-3-BME proteins (Cockrell et al., 2010; Ramser et al., 2010; Zhang et al., 1997), a 14-3-3ζ K49E mutant is predicted to diminish the interaction. Because the previously reported TPD systems have a high turnover rate (Clift et al., 2017; Nabet *et al*., 2018; Nishimura et al., 2009), we reasonably speculated that each individual difopein-EGFP protein that interacts with TDPP will also be transiently degraded and impossible to detect by co-immunoprecipitation (co-IP). Thus, instead of using a whole TDPP, we used only the bait part of TDPP, 14-3-3ζ (1-230) 20KR-HA, to evaluate its interaction with difopein-EGFP, along with the K49E mutant, 14-3-3ζ (1-230) 19KR K49E-HA, as the control. While although K49 has been mutated to R in the 14-3-3ζ (1-230) 20KR-HA construct, the interaction of 14-3-3ζ (1-230) 20KR-HA with difopein-EGFP can be detected by Immunoprecipitation, K49E mutation abolishes the interaction of 14-3-3ζ (1-230) 20KR-HA with difopein-EGFP (Fig. 3F). Also, as expected, the single K49E mutation also significantly diminished the degradation efficiency (Fig. 3, G and H), indicating an interaction-dependent degradation of difopein-EGFP by TDPP.

### TDPP degrades P-14-3-3-BME proteins

Beyond the evaluator protein difopein-EGFP, P-14-3-3-BME proteins are the major POIs of TDPP. To assess the influence TDPP has on the general P-14-3-3-BME proteins, the overall phospho-(Ser)-14-3-3-binding-motif level of HEK293T/17 cells was measured with its specific antibody, which was selected to reflect the general P-14-3-3-BME proteins level, and TDPP was again expressed in cells by transfection. During the evaluation, okadaic acid (OA), an inhibitor for PP2a and a partial inhibitor for PP1 (Dounay and Forsyth, 2002), was also used to enhance the total phosphorylation level. As expected, the overall phospho-(Ser)-14-3-3-binding-motif level in cells decreases by introducing TDPP (Fig. 4, A-C). The decrease can be observed under both normal conditions (Fig. 4A, lane 1 vs. 3) and dephosphorylation-inhibited treatment (Fig. 4A, lane 2 vs. 4), although exceptions still exist. Overall, these data show that TDPP can efficiently degrade P-14-3-3-BME proteins under both normal and high-phosphorylation conditions.

**Figure 4.**
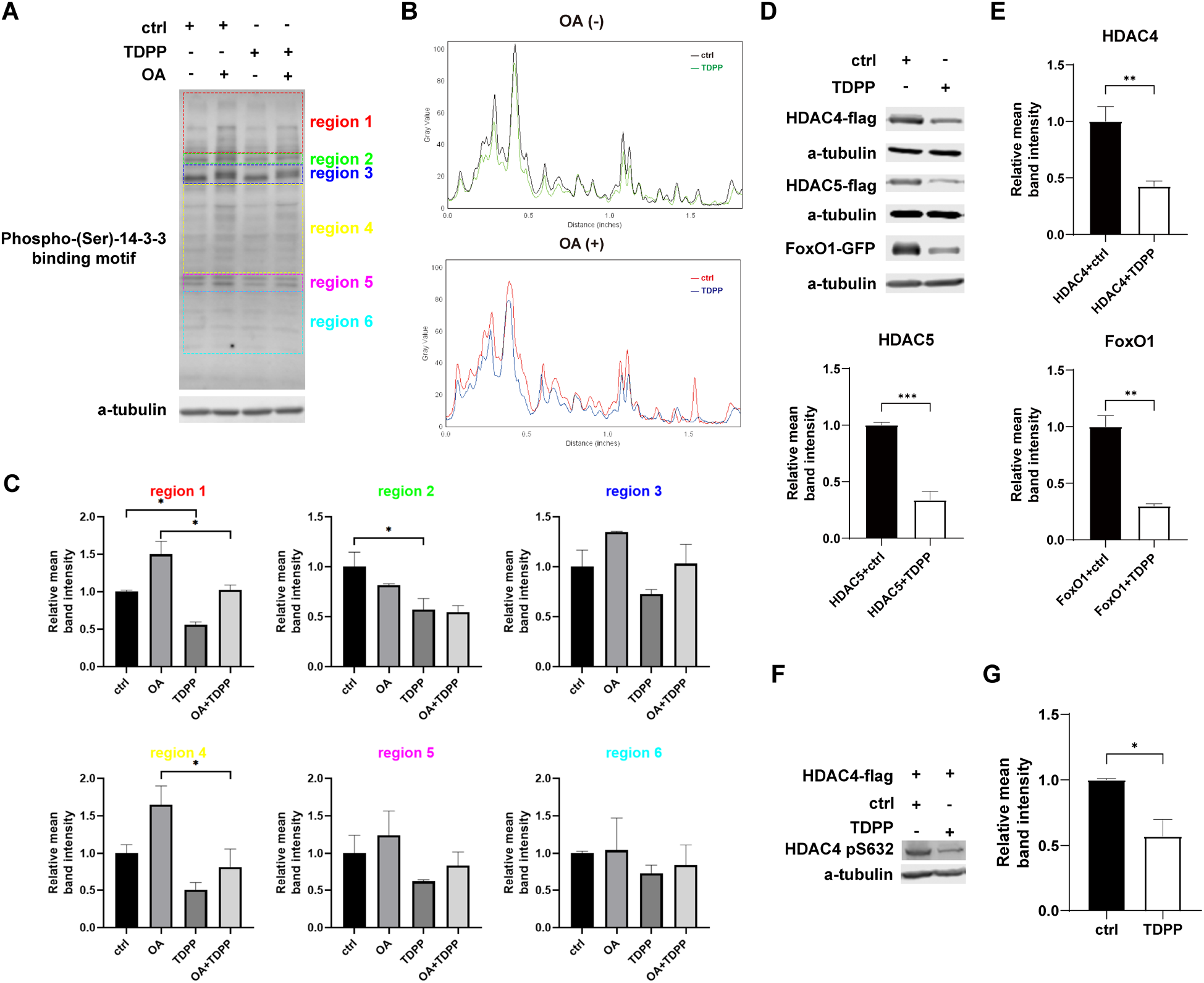
TDPP degraded phospho-14-3-3-binding-motif embedded proteins. A. Western blot of phospho-(Ser)-14-3-3-binding-motif treated with control/TDPP vector with/without 10 nM OA. B. Illustration of the grey value distribution of phospho-(Ser)-14-3-3-binding-motif in each treatment in panel A. X axis represents the distance to the top edge of the blot results. Upper panel: lane 1 (black) vs. lane 3 (green). Lower panel: lane 2 (red) vs. lane 4 (blue). C. Quantification of phospho-(Ser)-14-3-3-binding-motifembedded proteins with different molecular weight in panel A. Different molecular weight ranges were grouped as different regions and marked as different colors, which were roughly grouped by similar band intensity and reflect the difference among treatments. D and E. Western blot results (D) and the quantifications (E) of several different phospho-14-3-3-binding-motif embedded proteins (HDAC4, HDAC5, FoxO1) in HEK293T/17 cells treated with control/TDPP vectors. F and G. Western blot result (F) and the quantification (G) of HDAC4 phospho-S632 in a HDAC4 wildtype vectors transfected HEK293T/17 cells treated with control/TDPP vector.

To determine if TDPP’s degradation efficiency varies among different phosphoproteins, we further evaluated four randomly selected P-14-3-3-BME proteins. Three proteins showed significantly decreased protein levels by introducing TDPP (Fig. 4, D and E), while no significant difference was observed for CFL1 (Supplementary Fig. 2, A and B). We reasoned this variance may be due to the different 14-3-3-interaction ability and different subcellular distribution.

We next determined if the observed degradation is phosphorylation-dependent. A member of class IIa HDACs, histone deacetylase 4 (HDAC4), which plays an important role in tissue-specific growth and development (Wang et al., 2014), was selected as a known P-14-3-3-BME protein (Grozinger and Schreiber, 2000; Nishino et al., 2008; Paroni et al., 2008; Wang et al., 2000; Zhao et al., 2001) for further evaluation. The phospho-HDAC4 (pHDAC4)-14-3-3 interaction has been well-established (Kondo et al., 2019). HDAC4 phospho-S632 (pS632), which bears a P-14-3-3 binding motif, is a portion of HDAC4 that could phosphorylation-dependently interacted with 14-3-3. With an antibody that specifically detects HDAC4 pS632, we observed that TDPP can target HDAC4 pS632 (Fig. 4, F and G), consolidating a phosphorylation-induced degradation mechanism of TDPP. Taken together, these data established a phosphorylation-dependent degradation of P-14-3-3-BME proteins induced by TDPP.

### TDPP validates BICRA as a new 14-3-3 interaction target protein

Another application of TDPP is to validate potential P-14-3-3-BME proteins. We have previously worked on the SWI/SNF complex (Gao et al., 2008; Lei et al., 2013; Lei et al., 2019; Lei et al., 2015; Wang et al., 2004) that is involved in 14-3-3-related pathways (Parua et al., 2014). We thus applied TDPP to determine the potential agent that mediates 14-3-3-SWI/SNF interaction. Using 14-3-3-Pred (Madeira et al., 2015), we identified BICRA as a potential P-14-3-3-BME subunit of the SWI/SNF complex; the S801, S805, S815, S866, and S1477 residues in BICRA were predicted as potential 14-3-3-binding sites. The protein level of BICRA decreased substantially with the introduction of TDPP (Fig. 5, A and B), supporting BICRA as a P-14-3-3-BME subunit. This experiment suggests TDPP as a useful tool for validating potential 14-3-3 targeting proteins.

**Figure 5.**
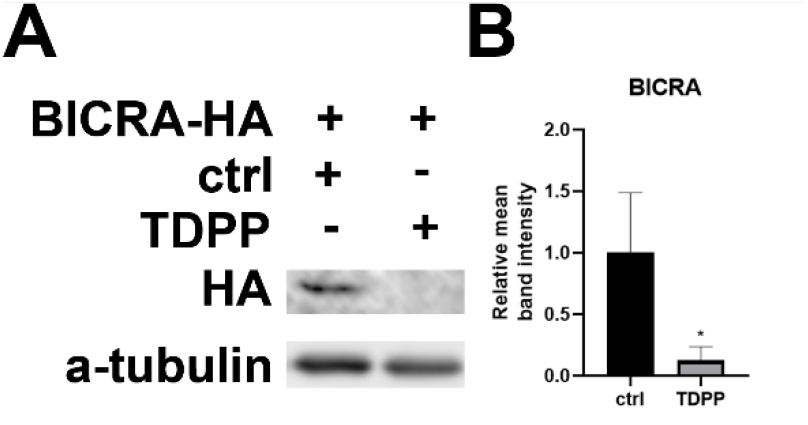
TDPP could predict potential 14-3-3 binding proteins. A. Western blot of BICRA-HA level in its expression HEK293T/17 cells transfected with control/TDPP vectors. B. Relative mean band intensity of BICRA-HA western blot.

## Discussion

In this study, we report for the first time a phosphorylation-induced TPD strategy, TDPP, which specifically degrades P-14-3-3-BME proteins by ligating an engineered 14-3-3ζ protein to a modified VHL E3-ligase. As designed, this protein chimera can efficiently bind and present P-14-3-3-BME proteins to E3-ligase which then introduces the targeted P-14-3-3-BME proteins to UPS for degradation (Fig. 1A).

Several innovations were applied in the design of TDPP’s protein structure. The most important one is the KR mutation to block the auto-ubiquitination and auto-degradation, which is predictable because of the protein nature of TDPP. One concern about the modification is that K49 is the key phosphoprotein-binding residue (Mortenson et al., 2015; Pennington et al., 2018; Zhang *et al*., 1997), by which the KR mutation would impair the interaction ability of TDPP with the POIs in this study. However, on one hand, it is also reported that multiple KR mutations in 14-3-3 could lead to a much more complex consequence to its interaction ability with P-14-3-3-BME proteins (Choudhary et al., 2009), which may counteract or reverse the effect of K49R mutation. On the other hand, arginine residue holds positive charge, just like the lysine residue, which represents that the KR mutation is almost like a synonymous substitution. Glutamic acid residue, on the opposite, holds negative charge, thus KE mutation could totally disrupt the core interaction ability of the mutated protein with its binding partners. In this study, the *in silico* prediction (Fig. 2) showed TDPP reserves the interaction ability with P-14-3-3-BME proteins, and the *in vivo* results (Fig. 3F) showed that significant interactions still exist between TDPP and our POIs.

The high efficiency of TDPP has been confirmed by a diversity of evaluators. Compared with P-14-3-3-BME proteins, the stable difopein-EGFP reporter, which does not have a phosphorylation/dephosphorylation switch and which has a weaker interaction with other unspecified proteins in cells, is appropriate for examining TDPP’s degradation efficiency (Fig. 3A). As expected, significant degradation of difopein-EGFP by TDPP was observed (Fig. 3, B-E) and, in contrast, the TDPP K49E significantly abolished degradation efficiency (Fig. 3, G and H), strongly indicating that TDPP is an interaction-based degradation system that specifically targets 14-3-3-binding proteins.

TDPP displays an overall high efficiency in degrading P-14-3-3-BME proteins (Fig. 4), though some variance in degradation efficiency was observed for different phosphoproteins. This is understandable because a few factors including the phosphorylation percentage of P-14-3-3-BME proteins, the half-life time, the spatiotemporal distribution, and the ability to bind 14-3-3 can all impact the apparent degradation efficiency. For instance, the binding affinity to 14-3-3 can have more than a 1,000-fold span among different P-14-3-3-BME proteins (Gogl et al., 2021). It is reasonable that TDPP may fail to degrade some proteins even if they bear a phsopho-14-3-3 binding motif, in which case TDPP could be locally saturated by POIs with relatively high binding affinity, while the low binding affinity targets could not be efficiently degraded..

Unlike that most of the previously reported TPD systems were designed for the degradation of one specific POI respectively, TDPP was designed to degrade the P-14-3-3-BME proteins, which are multiple proteins grouped together by a structure similarity and the ability to interact with 14-3-3 while phosphorylated. Thus, according to the nature of TDPP-induced degradation, “motif of interest (MOI)” or “PTM of interest (PTMOI)” will be better than the widely used “protein of interest” concept to describe the targets of TDPP, i.e., the “single” target of TDPP is P-14-3-3 binding motif. While P-14-3-3-BME proteins are mostly within the 14-3-3 interactome, such a strategy could be significant to investigate the P-14-3-3-BME proteins. The P-14-3-3-BME proteins are a group of proteins defined by a specific sequence within their protein structure. Though bearing special features, the P-14-3-3-BME proteins are not fully defined. It is challenging to determine the 14-3-3 binding motif with traditional methods like co-IP. Although computational tools have been developed to predict 14-3-3 interacting sites, a precise method to validate the predicted results is missing. Since TDPP specifically targets P-14-3-3-BME proteins, we believe that it will be a useful tool for validating potential P-14-3-3-BME proteins including BICRA studied here. Furthermore, TDPP can potentially target SARS-CoV-2, as it is reported that SARS-CoV-2 nucleocapsid protein is recognized by 14-3-3 (Tugaeva et al., 2021).

Protein phosphorylation/dephosphorylation plays as a “switch” to activate or inactivate phosphatidylinositol 3-kinase (Hemmings and Restuccia, 2012). A simple degradation of the total protein is unspecified and the degradation of proteins with or without PTMs cannot be told apart. PPIs are also affected by PTMs (Keskin et al., 2016). Proteins with PTMs can be recognized by their specific “readers”, such as Forkhead-associated (FHA) domains for protein phosphorylation (Almawi et al., 2017), and bromodomains for histone acetylation (Fujisawa and Filippakopoulos, 2017). For P-14-3-3-BME proteins, three kinds of the binding motifs have been identified, RSXpSXP (mode I), RXXXpSXP (mode II), and (pS/pT)X_0-2_-COOH (mode III), where pS/pT represents phosphoserine/phosphothreonine (Garnier et al., 2021; Yang et al., 2006). Based on the design and our results, it is presumed that TDPP could have great degradation efficiency for proteins with those three kinds of motifs. While 14-3-3 and P-14-3-3-BME proteins are involved in many cellular events, including neural development (Antunes et al., 2022), cancer proliferation and apoptosis (Aljabal and Yap, 2020; Huang et al., 2020; Yang et al., 2020), autoimmune disorder (McGowan et al., 2020), virus infection (Liu et al., 2021), etc., the TDPP-induced degradation could be significant in regulating the related downstream pathways and biological events.

The strategy of designing TDPP is scalable and it can be easily extended to target other proteins that share similar characteristics. For example, H3K9ac-embedded histones may be targeted by bromodomain-embedded proteins ligated with E3-ligases. We are enthusiastic that the design of TDPP will inspire more targeted degradation designs, especially those targeting a specific interactome, and thus inspiring more ideas in the designing of TPD systems.

One concern about TDPP is its targeting specificity. Certain potential mechanisms may make TDPP-induced degradation off-target. In theory, some non-P-14-3-3-BME proteins may be degraded by TDPP through nonspecific binding to 14-3-3 (i.e., not bind to the binding groove of 14-3-3). Specifically, one scenario that we should pay more attention to is the 14-3-3 dimerization (Shen et al., 2003), where the heterodimerization of TDPP and an endogenous 14-3-3 protein could potentially deplete 14-3-3 in cells. Another case is condensates existing in cells (Lyon et al., 2021); some proteins may condense to a specific cellular location and have specific functions. Such a condensate behavior, however, may lead TDPP to target a protein without a phospho-14-3-3 binding motif. Therefore, TDPP as a prototype for phosphorylation-based degradation system will need to be customized for further specific application.

In summary, we describe a TPD strategy, the TDPP system, that specifically targets and degrades P-14-3-3-BME proteins. We have validated the degradation efficiency and mechanism of TDPP with a difopein-EGFP system, and general and specific P-14-3-3-BME proteins. As the first phosphorylation-induced TPD system, TDPP not only can inspire more similar strategies in TPD designing, but also could be a powerful tool to regulate phospho-14-3-3 binding motif levels and validate unidentified 14-3-3 binding proteins, which are very important to 14-3-3 related research.

## Methods

### Plasmid information

Plasmids encoded with VHL (#19234), 14-3-3ζ (#1942), HDAC4 (#30485), HDAC4 3SA (#30486), HDAC5 (#32213), FoxO1 (#9022), CFL1 (#78295), BICRA (#34902) were obtained from Addgene. Difopein DNA was constructed with 3 ssDNA oligos (sequence can be found in Supplement File 2) by overlapping PCR. Plasmids encoded with VHL 3KR-14-3-3ζ (1-230) 20KR mutation, VHL 3KR-14-3-3ζ (1-230) 19KR K49E mutation were synthesized by GenScript Biotech. The transfection vectors of each gene described above were constructed into pcDNA3.1 backbone using PCR amplification.

### Transfection

Transfection of HEK293T/17 cells was performed using Lipofectamine 2000 (Thermo Fisher Scientific) according to manufacturer’s protocol.

### Chemical information

Okadaic Acid (#10011490), MLN7243 (#30108), and MG132 (#10012628) were purchased from CAYMAN Chemical.

### Antibodies

The antibodies used in this study were as follows: Antibodies targeting α-tubulin (#2125 and #3873), Phospho-(Ser) 14-3-3 Binding Motif Antibody (#9601), HA-Tag (#2367) were purchased from Cell Signaling Technology. Antibodies targeting EGFP (ab290), V5 (ab9116) were purchased from Abcam. ANTI FLAG antibody (F1804) was purchased from Sigma.

### Mammalian cells and cell culture

HEK 293T/17 was purchased from ATCC (CRL-11268). Cells were cultured according to ATCC’s protocol with minor optimizations. For a typical cell culture cycle with transfection, cells were seeded at 50×10^3^ cells/cm^2^ density, and 24 hours after cell seeding, attached cells were transfected with plasmid or plasmids mix encoded with desired gene(s) with Lipofectamine 2000 following manufacture’s protocol. 48 hours after transfection, cells were collected for further evaluations.

### Western blotting

Adherent cells were washed with pre-cooled Hank’s balanced salt solution (HBSS, Gibco #14025092) supplemented with phosphatase inhibitors (Sigma #4906845001) and protease inhibitor cocktail (Sigma #11873580001). Cell pellets or protein supernatants were lysed in 1x Laemmli sample buffer containing beta-mercaptoethanol, phosphatase inhibitors and protease inhibitor cocktail. The whole-cell lysates were normalized to same protein concentration and then separated using SDS-PAGE, sequenced by transferring onto polyvinyllidene difluoride membranes. The membrane was incubated with a blocking solution consisting of3% bovine serum albumin (BSA) in TBS solution for phosphorylated proteins of interest, or 4% non-fat dry milk (NFDM) in PBS solution for other proteins of interest. The membrane was blocked for 3 times and 10 min each time at room temperature, then incubated with primary antibodies in blocking solution overnight at 4°C, followed by incubation with host-specific Alexa Fluor-conjugated or HRP-conjugated secondary antibodies (1:10000 dilution) for 1h. For signal detection, the membrane was developed directly with a LI-COR Imaging System for Alexa Fluor-conjugated secondary antibodies or with a mixture of ECL solution (Thermo Fisher Scientific) using X-ray films. The developed bands were analyzed with Image Studio (LI-COR), Image Lab (Bio-Rad), or ImageJ (NIH) software. All unedited full-length gel images could be found in Supplement File 3.

### Co-immunoprecipitation

The co-immunoprecipitation method was performed to investigate protein-protein interactions. HEK293T/17 cells were transfected with indicated constructs as described in “Transfection” section. Adherent cells were washed with cold HBSS supplemented with phosphatase inhibitors and protease inhibitor cocktail, sequenced by cells scraping and lysing the cell pellets in immunoprecipitation lysis buffer (150 mM NaCl, 1% Triton X-100 in 20 mM HEPES, supplemented with phosphatase inhibitors and protease inhibitor cocktail) for 30 min on rotator at 4 °C. Next, the supernatant was separated by centrifuging at 13,000 g at 4 °C for 5 min and collected, to which the normal mouse IgG (Santa Cruz sc2025), normal rabbit IgG (Cell Signaling #2729), or antibodies against protein of interest, and Protein G Dyna beads (Fisher #10004D) were supplemented and incubated with the cell lysate at 4 °C on a rotor overnight. The beads were washed four times with IP buffer (150 mM NaCl, 0.05% TWEENS 20 in 20 mM HEPES, supplemented with phosphatase inhibitors and protease inhibitor cocktail), resuspended in 1x Laemmli Sample Buffer containing beta-mercaptoethanol, phosphatase inhibitors and protease inhibitor cocktail, followed by SDS-PAGE and immunoblotting with specified antibodies.

### Fluorescence assay

HEK293T/17 cells were transfected with indicated constructs as described in “Transfection” section. For difopein-EGFP transfected HEK293T/17 cells, the nuclei were stained with Hoechst 33342 (Fisher H3570) for 1 hr prior the fluorescence assay, and EGFP fluorescence assay was performed under an inverted fluorescence microscope (Olympus IX73P2F), followed by image analysis with ImageJ (NIH) software. For each microscope field, the fluorescence intensity under green and blue channel were considered as representative of EGFP intensity and cell number, respectively. The EGFP intensity was then normalized for each field by dividing intensity of green channel with intensity of blue channel.

### RNA isolation and quantitative real-time PCR

RNA was prepared by TRIzol Reagent (Invitrogen #15596018) following manufacturer’s instructions. Diluted RNA was reverse transcribed using iScript cDNA Synthesis Kit (Bio-Rad #1708891) following manufacturer’s instructions. Quantification of the gene expression was carried out by quantitative real-time PCR using the SYBR Green PCR Master Mix (Fisher #43-687-08) following manufacturer’s instructions. GAPDH was used as internal control. The primers are provided in Supplement File 2.

### Statistical analysis

Results were presented as mean ± S.E. Statistically significant differences between groups were analyzed by two-tailed, unpaired Student’s t-test or one-way analysis of variance (ANOVA) followed by the Student-Newman-Keuls multiple comparisons tests using Prism 8 (GraphPad) software. A p-value < 0.05 was regarded as significant. Each experiment was performed at least three times.

## Data availability

Data generated in this study are provided in the article and its associated files. Source data are provided with this paper. All other data are available from the corresponding authors on request.

## Acknowledgements

This research was supported in part by R01HL139735 and R01HL163672 to ZW. We would like to thank Dr. Shaomeng Wang for suggestions and Dr. Leonid Bnatovskiy for editing the manuscript.

Figure 1 and Figure 3A were created with BioRender.com.

## Author contributions

Z.L., E.C., Z.W. and L.L., conceptualization; Z.L., X.H. and L.L., data curation; Z.L., and L.L., formal analysis; Z.L., and L.L., validation; Z.L., X.H. and M.L., visualization; Z.L., writing-original draft; Z.L., X.H. and M.L., software; Z.L., X.H., Z.W. and L.L., methodology; Z.L., E.C., Z.W. and L.L., resources; E.C., Z.W. and L.L., investigation; E.C., Z.W. and L.L., project administration; X.H., Z.S., Z.W. and L.L., editing; E.C., Z.W. and L.L., supervision; Z.W., funding acquisition.

