## Supplement File 1 for "A ubiquitination-mediated degradation system to target phospho-14-3-3-binding-motif embedded proteins"

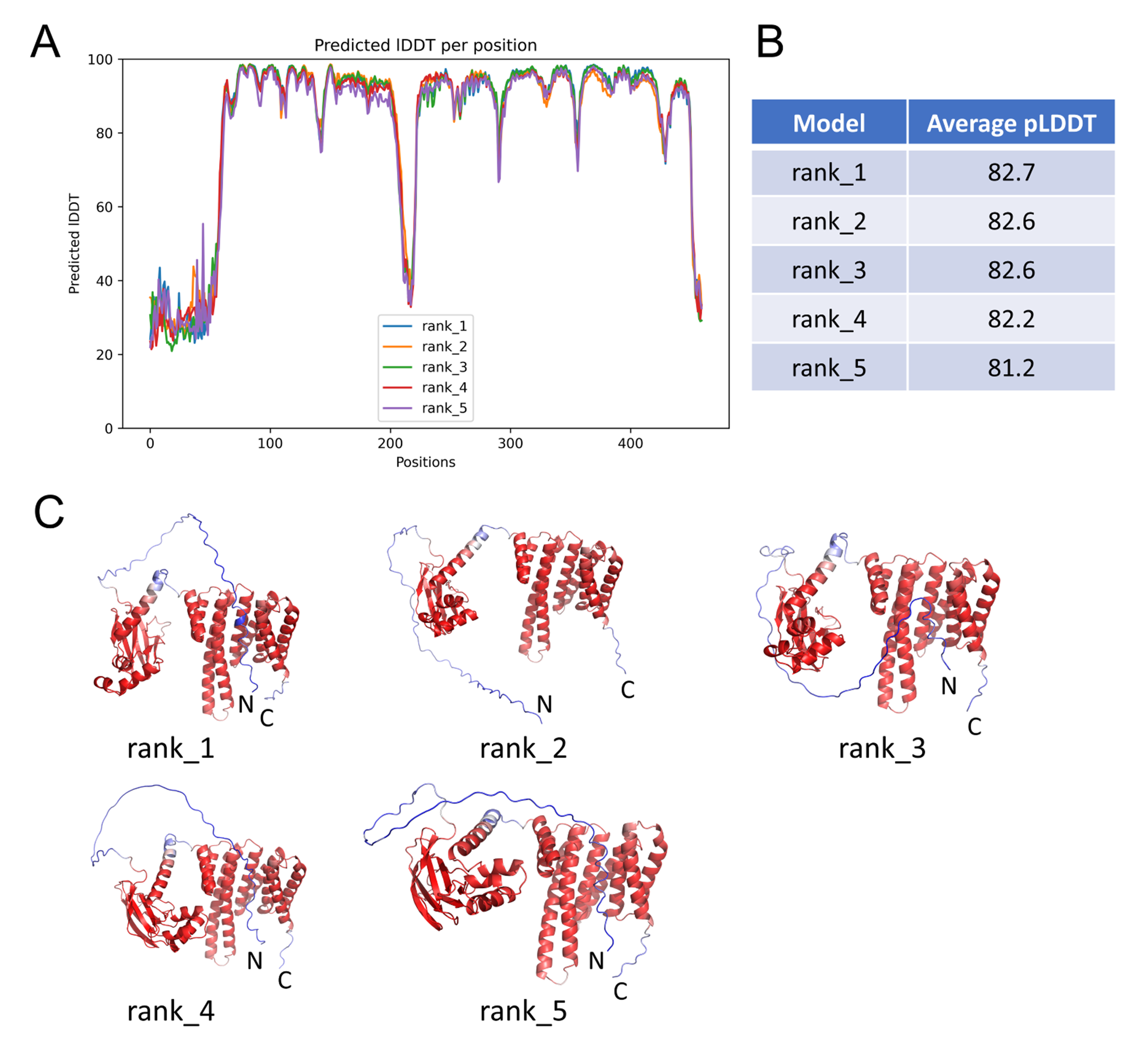
**Supplementary Figure 1.** A. Illustration of predicted per-residue local-distance difference test (LDDT) score for each model. B. Average pLDDT score. C. Illustration of five full-length models colored by pLDDT score (red: high, blue: low, and white: intermediate).

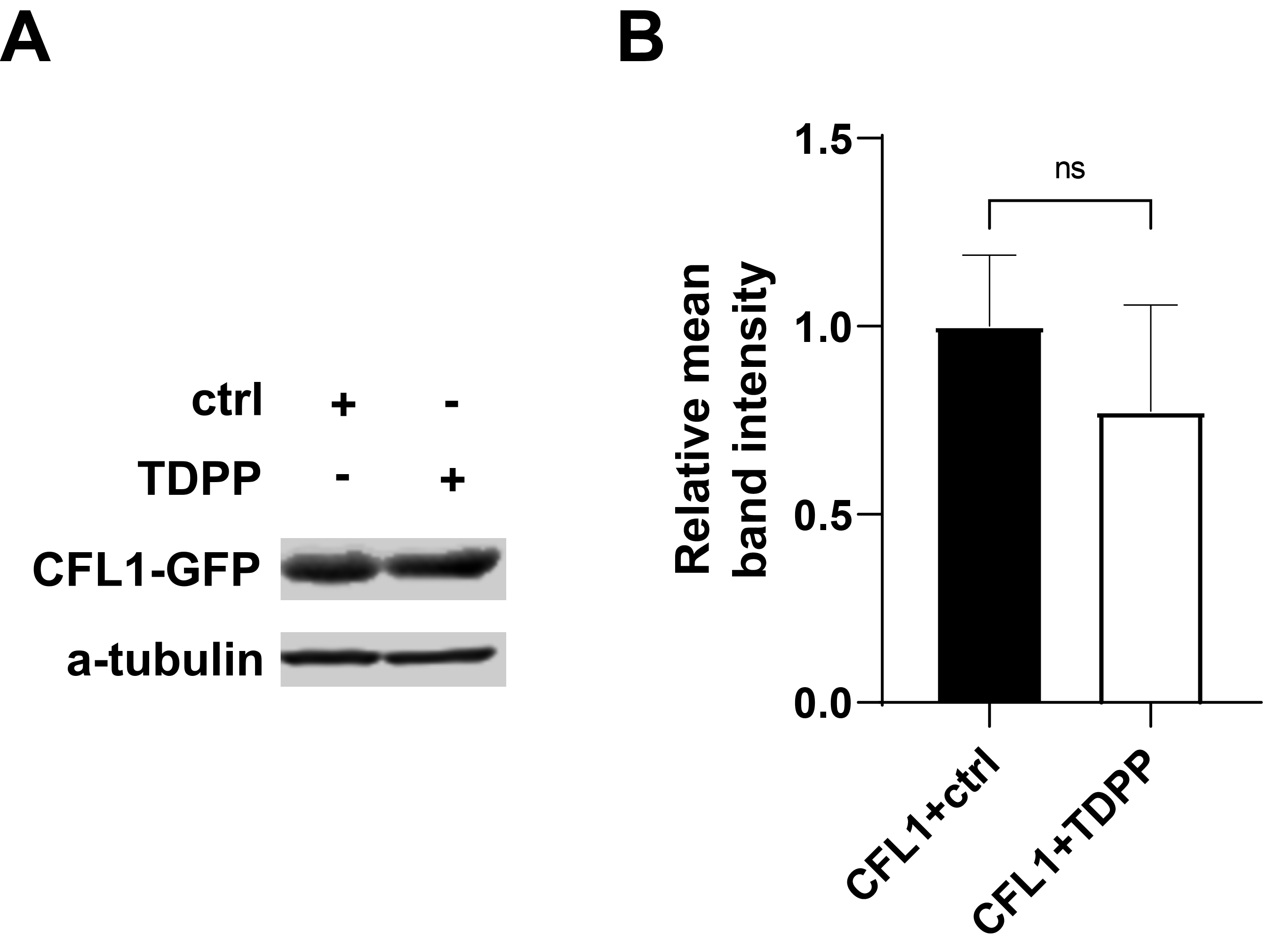
**Supplementary Figure 2.** Western blot results (A) and the quantification (B) of CFL1 in HEK293T/17 cells treated with control/TDPP vectors. Protein level of CFL1-GFP and are not significantly influenced by TDPP treatment.
