## Supplement File 2 for "A ubiquitination-mediated degradation system to target phospho-14-3-3-binding-motif embedded proteins"

DNA oligos and primers

1. Oligos used for difopein synthesis

Oligo 1: CCCACGCGTGCCACCATGTCTGCCGATGGTGCCCCGCACTGTGTTCCGCGTGATCTGTCTTGGCTGGATCTGGAAGCCAATATGTGTC

Oligo 2: CTCAGATCGCGCGGCACGCAATGCGGCGCGCCATCGGCAGAATCCAGACCGGCGGCACCCGGCAGACACATATTGGCTTCCAGA

Oligo 3: CCCGAATTCTTATTCCAGGCCCGCCGCGCCCGGCAGGCACATGTTCGCTTCCAGATCCAGCCAGCTCAGATCGCGCGGCACGCA

2. Primers used for overlapping PCR for difopein synthesis:

Primer 1: CCCACGCGTGCCACCATGTCTGCCGATGGTGCCCCGCA

Primer 2: CTCAGATCGCGCGGCACGCA

Primer 3: ACTGCCGCTGCCAGAGCCTTCCAGGCCCGCCGCGCCCG

3. Primers used for quantitative real-time PCR:

EGFP qPCR Forward: AGTCCGCCCTGAGCAAAGA

EGFP qPCR Reverse: TCCAGCAGGACCATGTGATC

GAPDH qPCR Forward: GTCTCCTCTGACTTCAACAGCG

GAPDH qPCR Reverse: ACCACCCTGTTGCTGTAGCCAA
