## Supplement File 3 for "A ubiquitination-mediated degradation system to target phospho-14-3-3-binding-motif embedded proteins"

### Slide 1
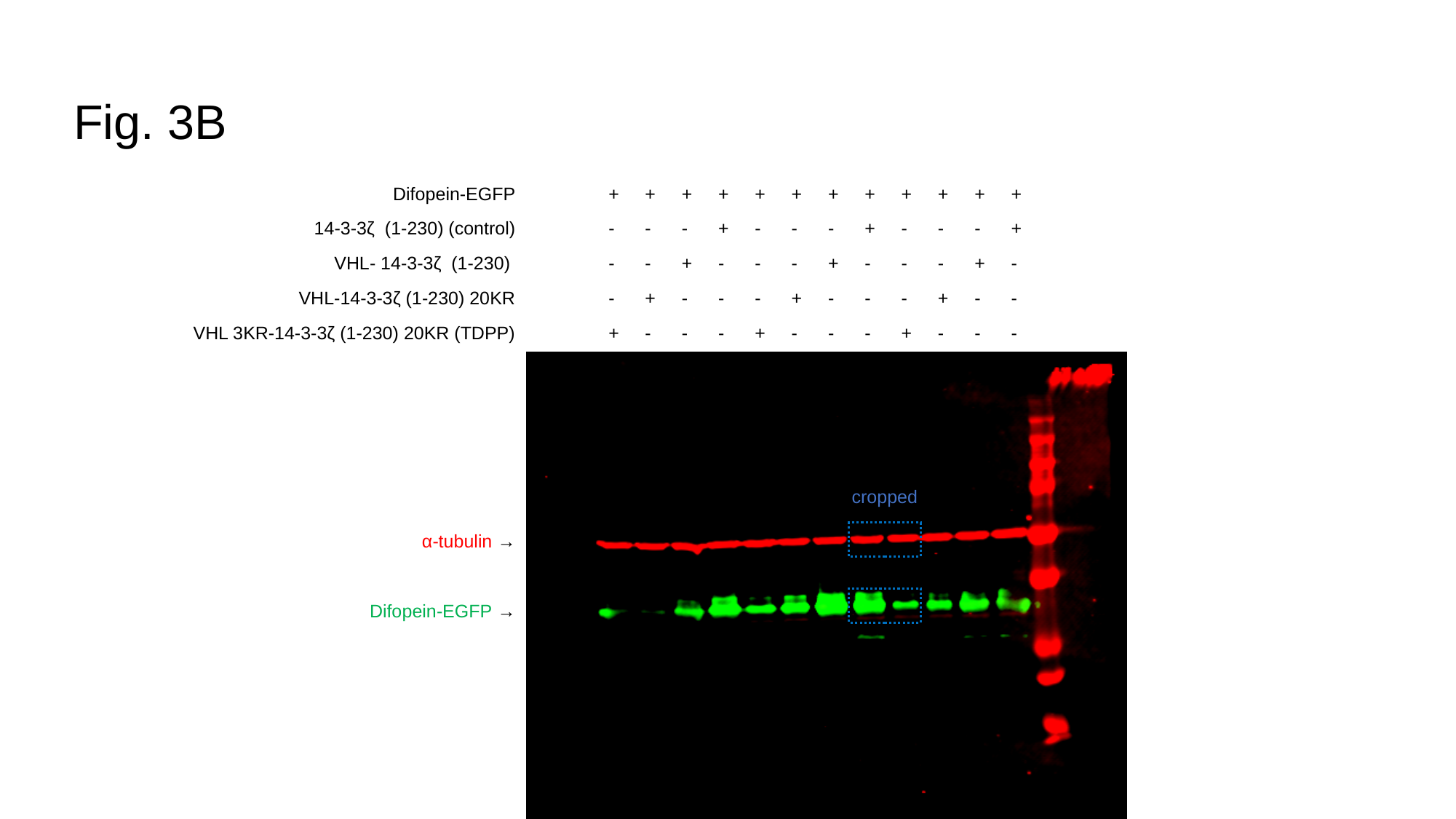

#### Fig. 3B
Difopein-EGFP
14-3-3ζ (1-230) (control)
VHL- 14-3-3ζ (1-230)
VHL-14-3-3ζ (1-230) 20KR
VHL 3KR-14-3-3ζ (1-230) 20KR (TDPP)
α-tubulin →
Difopein-EGFP →
+
-
-
-
+
+
-
-
+
-
+
-
+
-
-
+
+
-
-
-
+
-
-
-
+
+
-
-
+
-
+
-
+
-
-
+
+
-
-
-
+
-
-
-
+
+
-
-
+
-
+
-
+
-
-
+
+
-
-
-
cropped

### Slide 2
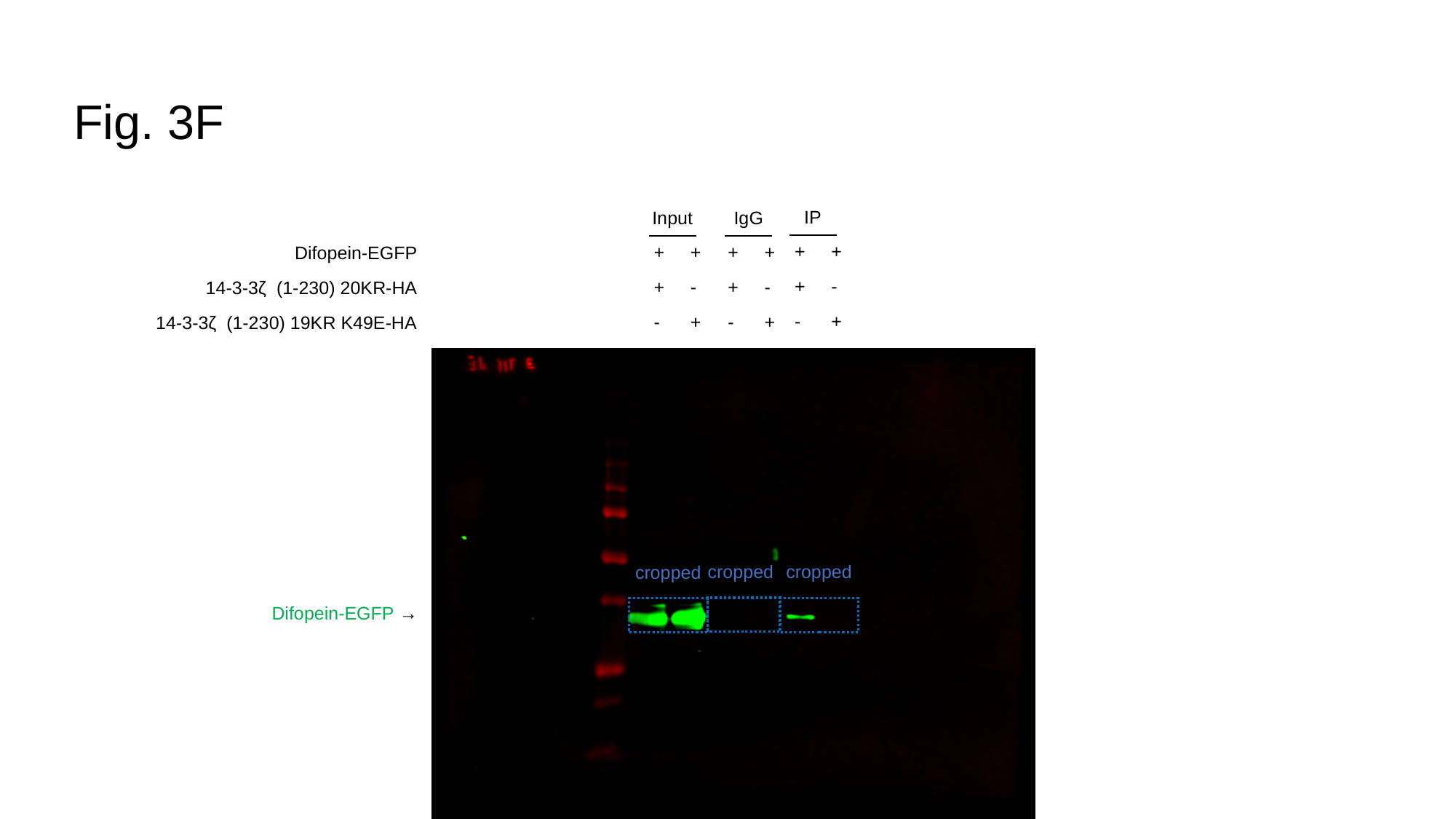

#### Fig. 3F
IP
Input
IgG
+
+
-
+
-
+
+
+
-
+
-
+
+
+
-
+
-
+
Difopein-EGFP
14-3-3ζ (1-230) 20KR-HA
14-3-3ζ (1-230) 19KR K49E-HA
cropped
cropped
cropped
Difopein-EGFP →

### Slide 3
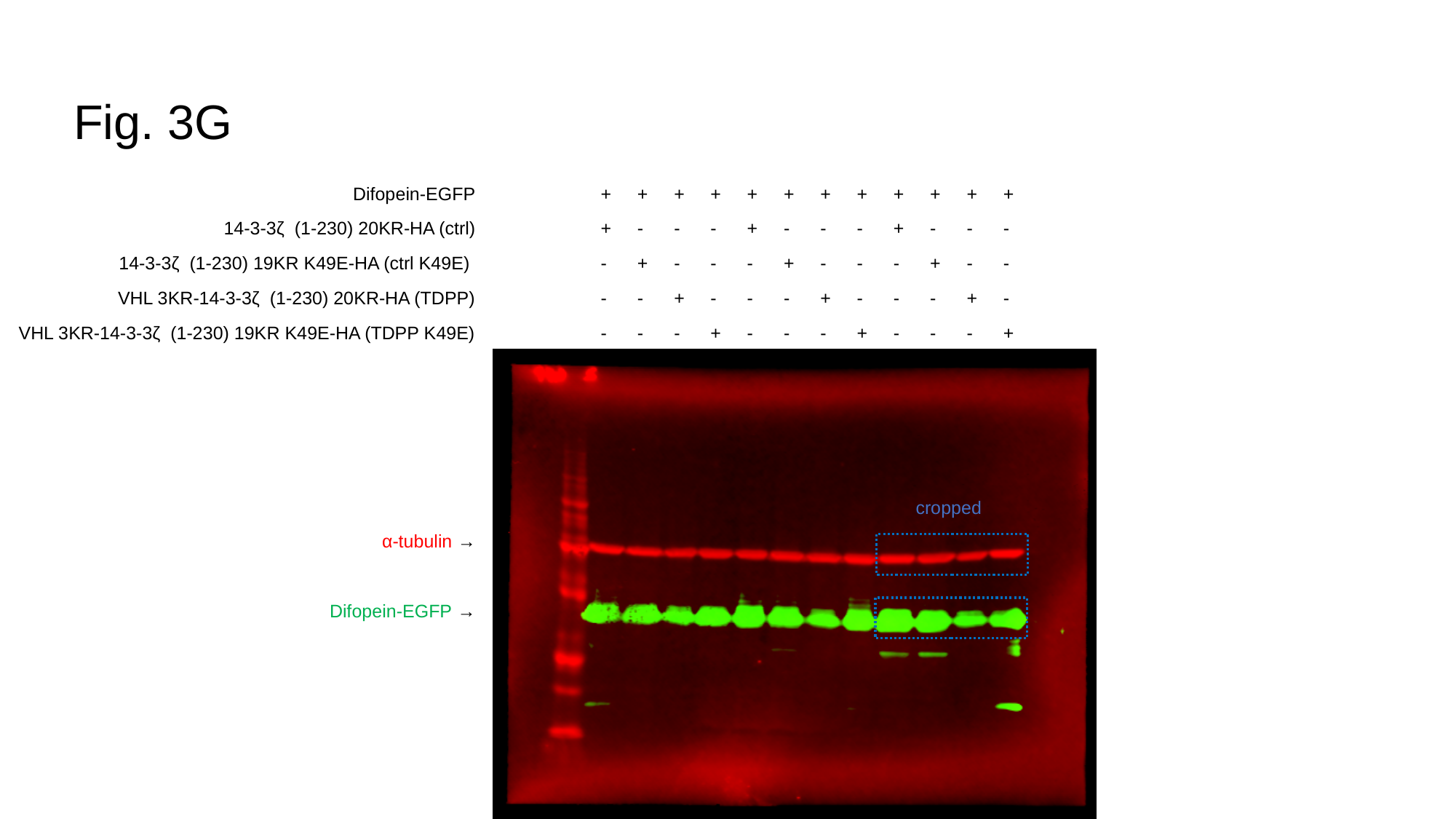

#### Fig. 3G
Difopein-EGFP
14-3-3ζ (1-230) 20KR-HA (ctrl)
14-3-3ζ (1-230) 19KR K49E-HA (ctrl K49E)
VHL 3KR-14-3-3ζ (1-230) 20KR-HA (TDPP)
VHL 3KR-14-3-3ζ (1-230) 19KR K49E-HA (TDPP K49E)
α-tubulin →
Difopein-EGFP →
+
+
-
-
-
+
-
+
-
-
+
-
-
+
-
+
-
-
-
+
+
+
-
-
-
+
-
+
-
-
+
-
-
+
-
+
-
-
-
+
+
+
-
-
-
+
-
+
-
-
+
-
-
+
-
+
-
-
-
+
cropped

### Slide 4
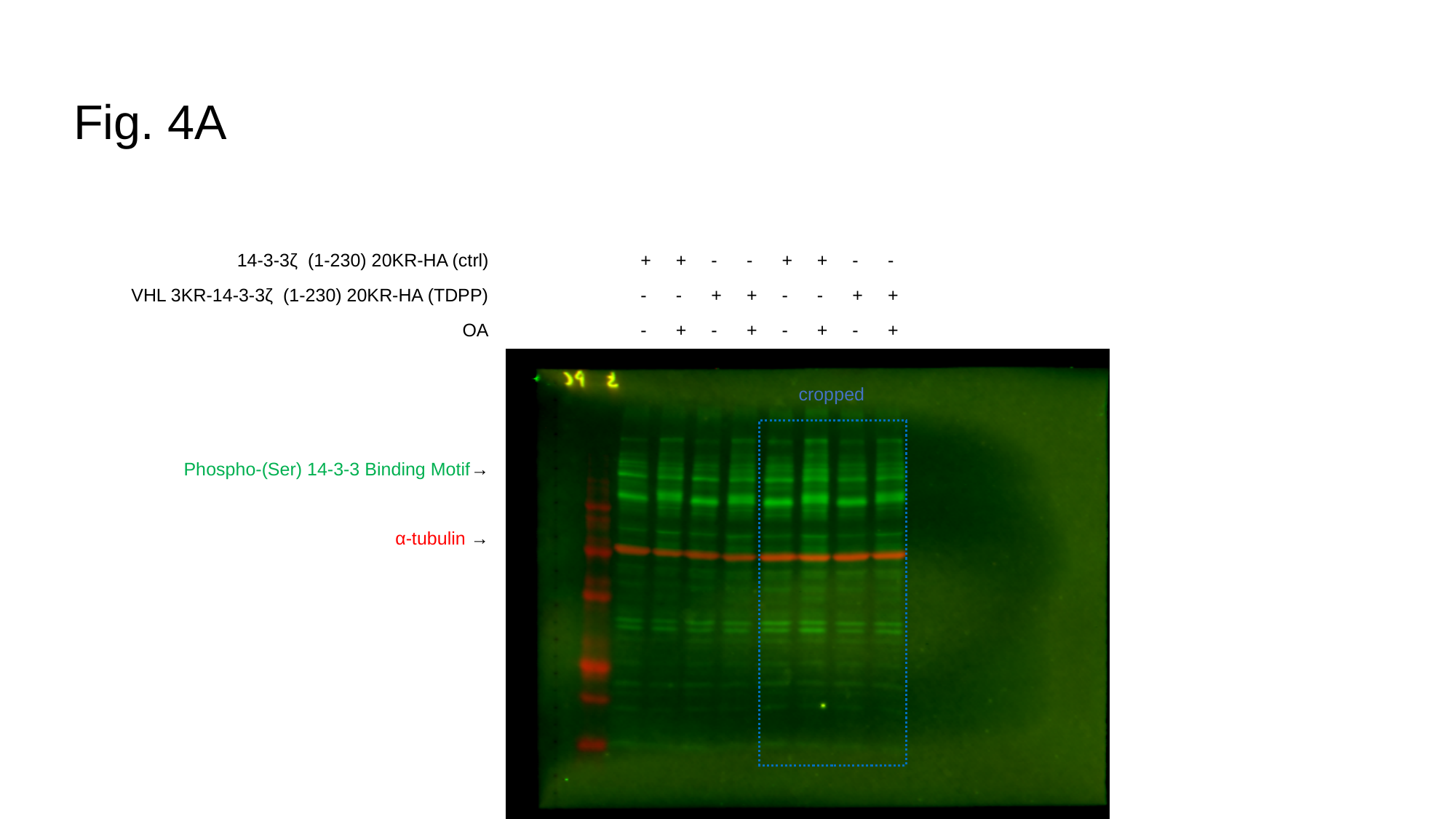

#### Fig. 4A
14-3-3ζ (1-230) 20KR-HA (ctrl)
VHL 3KR-14-3-3ζ (1-230) 20KR-HA (TDPP)
OA
Phospho-(Ser) 14-3-3 Binding Motif→
α-tubulin →
+
-
-
+
-
+
-
+
-
-
+
+
+
-
-
+
-
+
-
+
-
-
+
+
cropped

### Slide 5
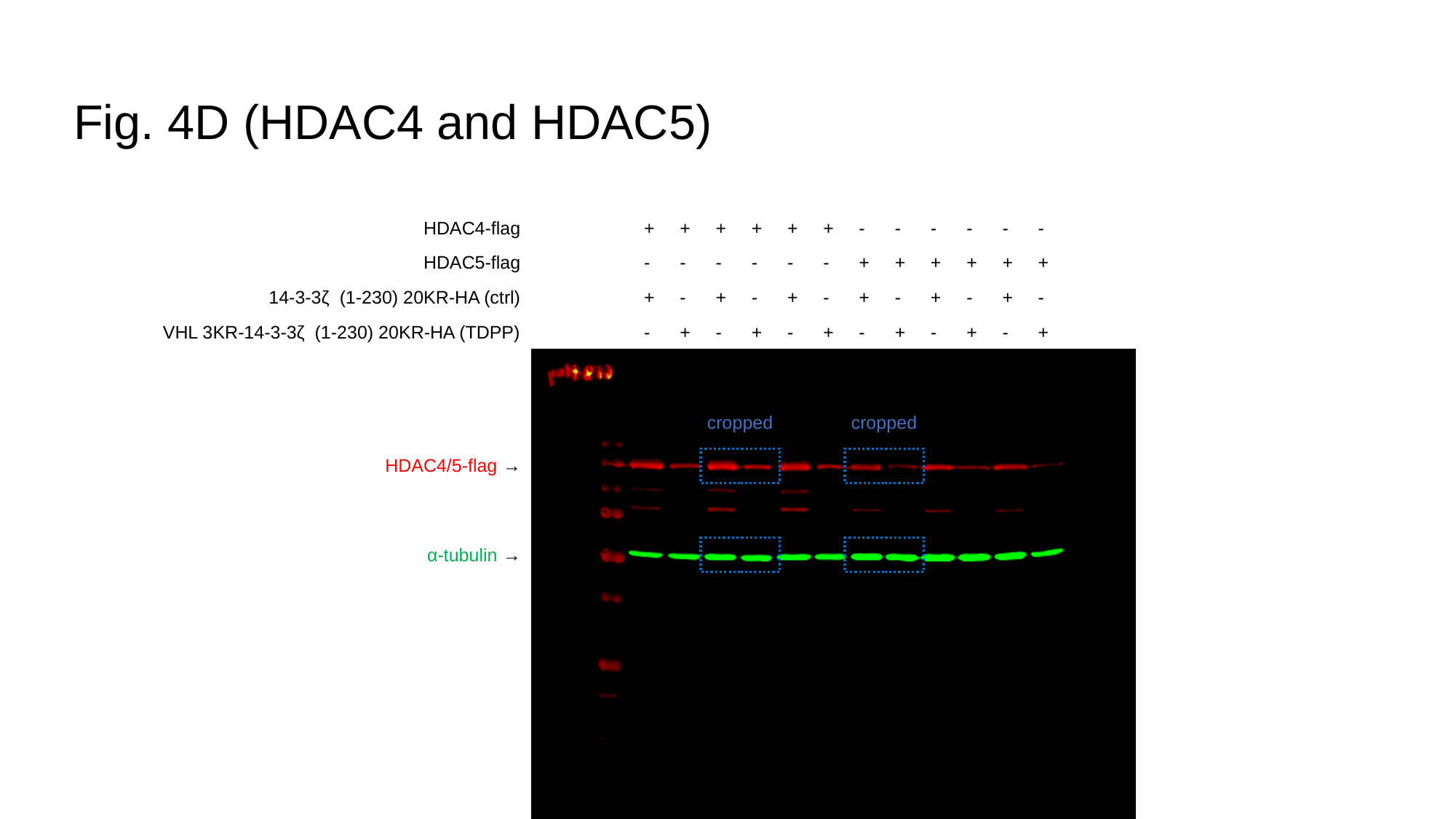

#### Fig. 4D (HDAC4 and HDAC5)
HDAC4-flag
HDAC5-flag
14-3-3ζ (1-230) 20KR-HA (ctrl)
VHL 3KR-14-3-3ζ (1-230) 20KR-HA (TDPP)
+
-
+
-
+
-
-
+
+
-
+
-
+
-
-
+
+
-
+
-
+
-
-
+
-
+
+
-
-
+
-
+
-
+
+
-
-
+
-
+
-
+
+
-
-
+
-
+
cropped
cropped
HDAC4/5-flag →
α-tubulin →

### Slide 6
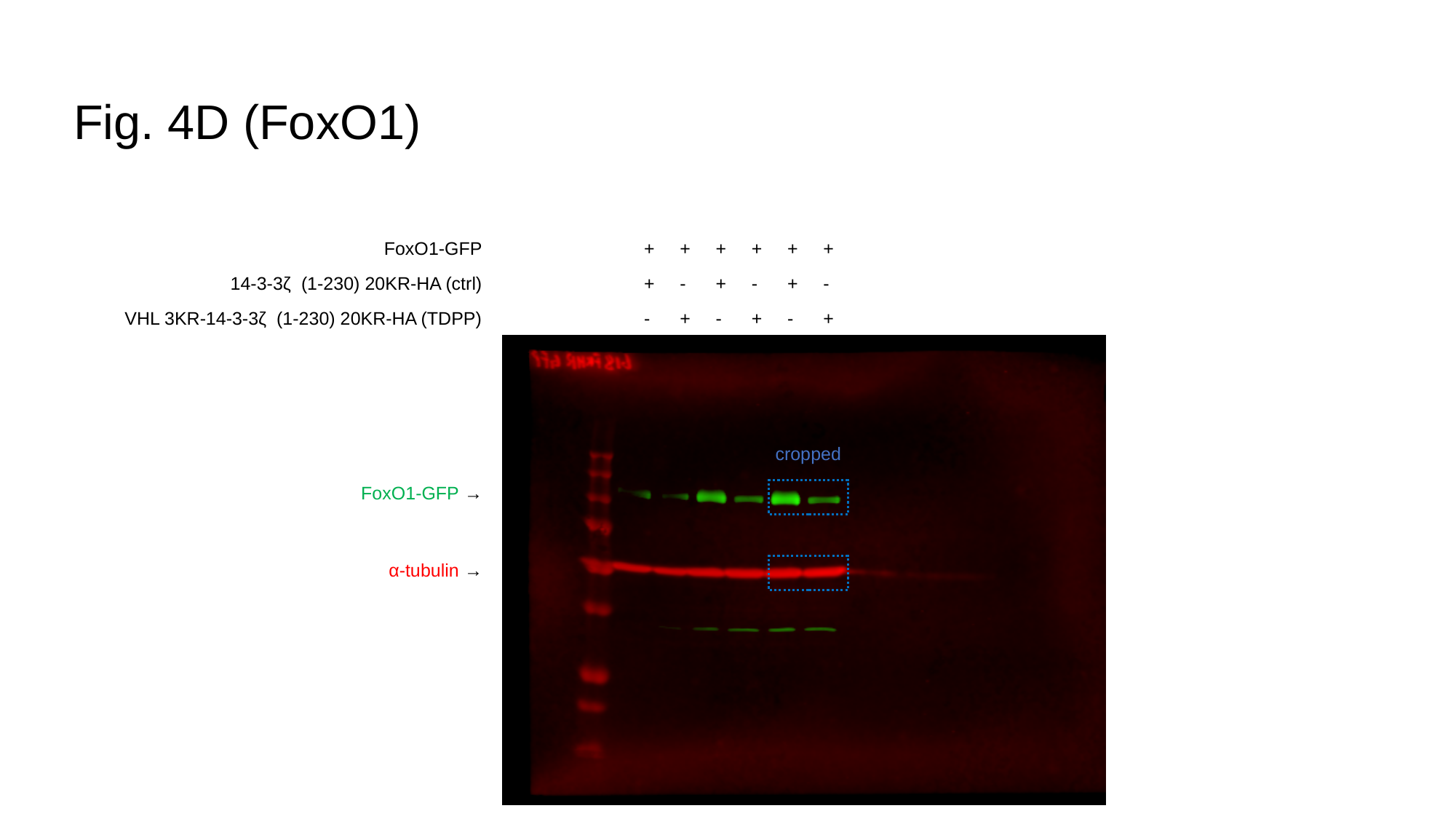

#### Fig. 4D (FoxO1)
FoxO1-GFP
14-3-3ζ (1-230) 20KR-HA (ctrl)
VHL 3KR-14-3-3ζ (1-230) 20KR-HA (TDPP)
+
+
-
+
-
+
+
+
-
+
-
+
+
+
-
+
-
+
cropped
FoxO1-GFP →
α-tubulin →

### Slide 7
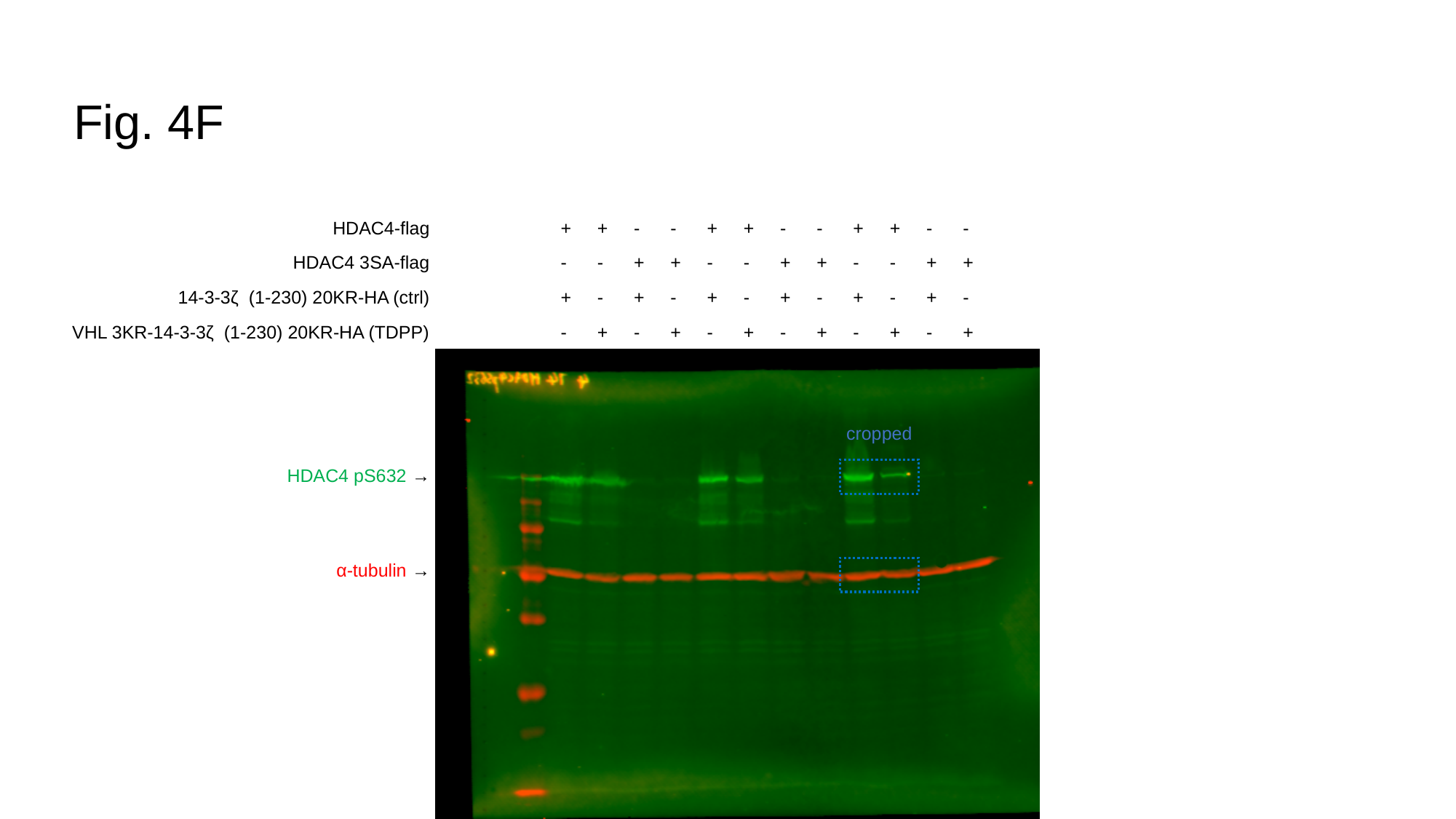

#### Fig. 4F
HDAC4-flag
HDAC4 3SA-flag
14-3-3ζ (1-230) 20KR-HA (ctrl)
VHL 3KR-14-3-3ζ (1-230) 20KR-HA (TDPP)
+
-
+
-
+
-
-
+
-
+
+
-
-
+
-
+
+
-
+
-
+
-
-
+
-
+
+
-
-
+
-
+
+
-
+
-
+
-
-
+
-
+
+
-
-
+
-
+
cropped
HDAC4 pS632 →
α-tubulin →

### Slide 8
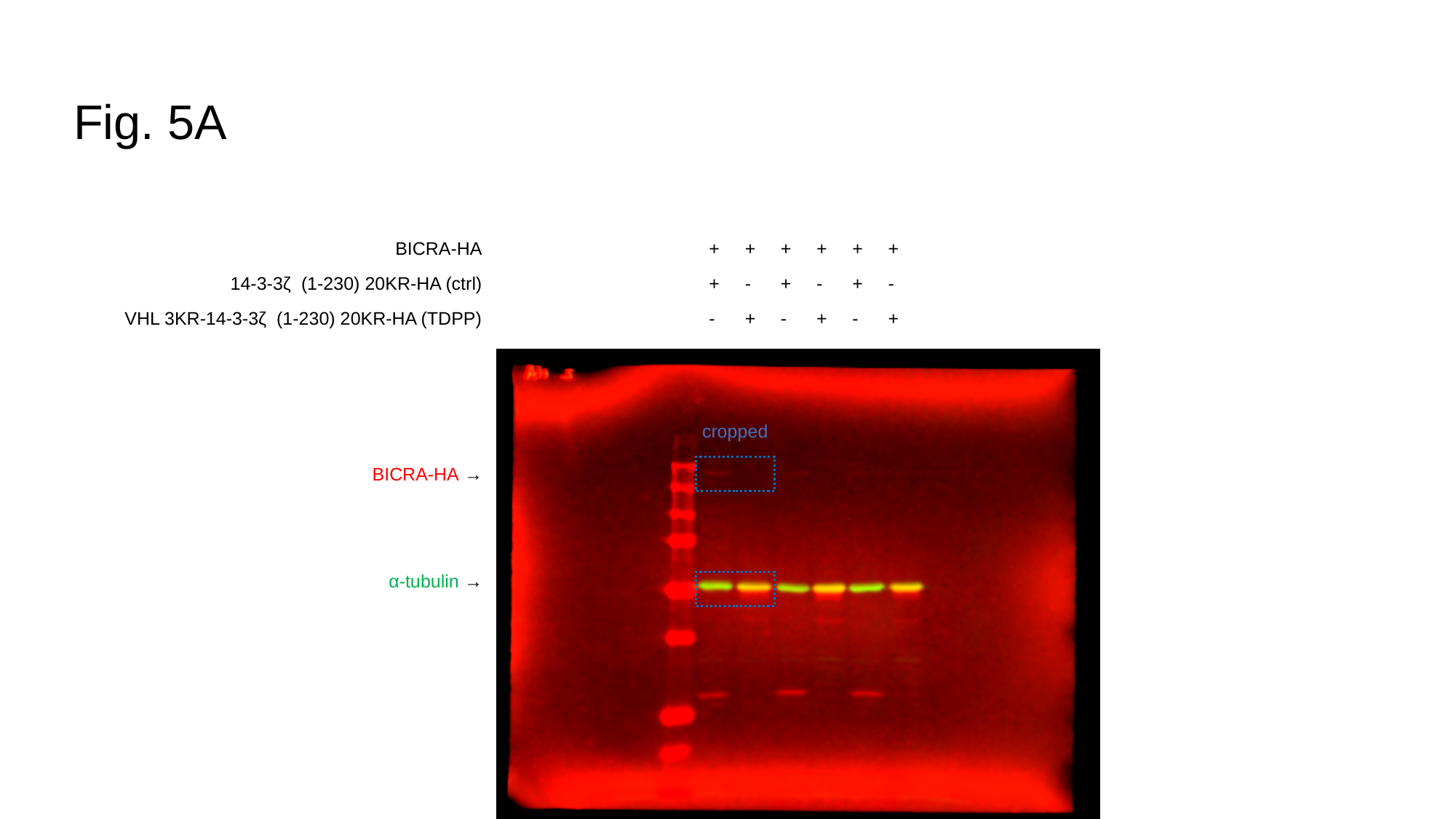

#### Fig. 5A
BICRA-HA
14-3-3ζ (1-230) 20KR-HA (ctrl)
VHL 3KR-14-3-3ζ (1-230) 20KR-HA (TDPP)
+
+
-
+
-
+
+
+
-
+
-
+
+
+
-
+
-
+
cropped
BICRA-HA →
α-tubulin →

### Slide 9
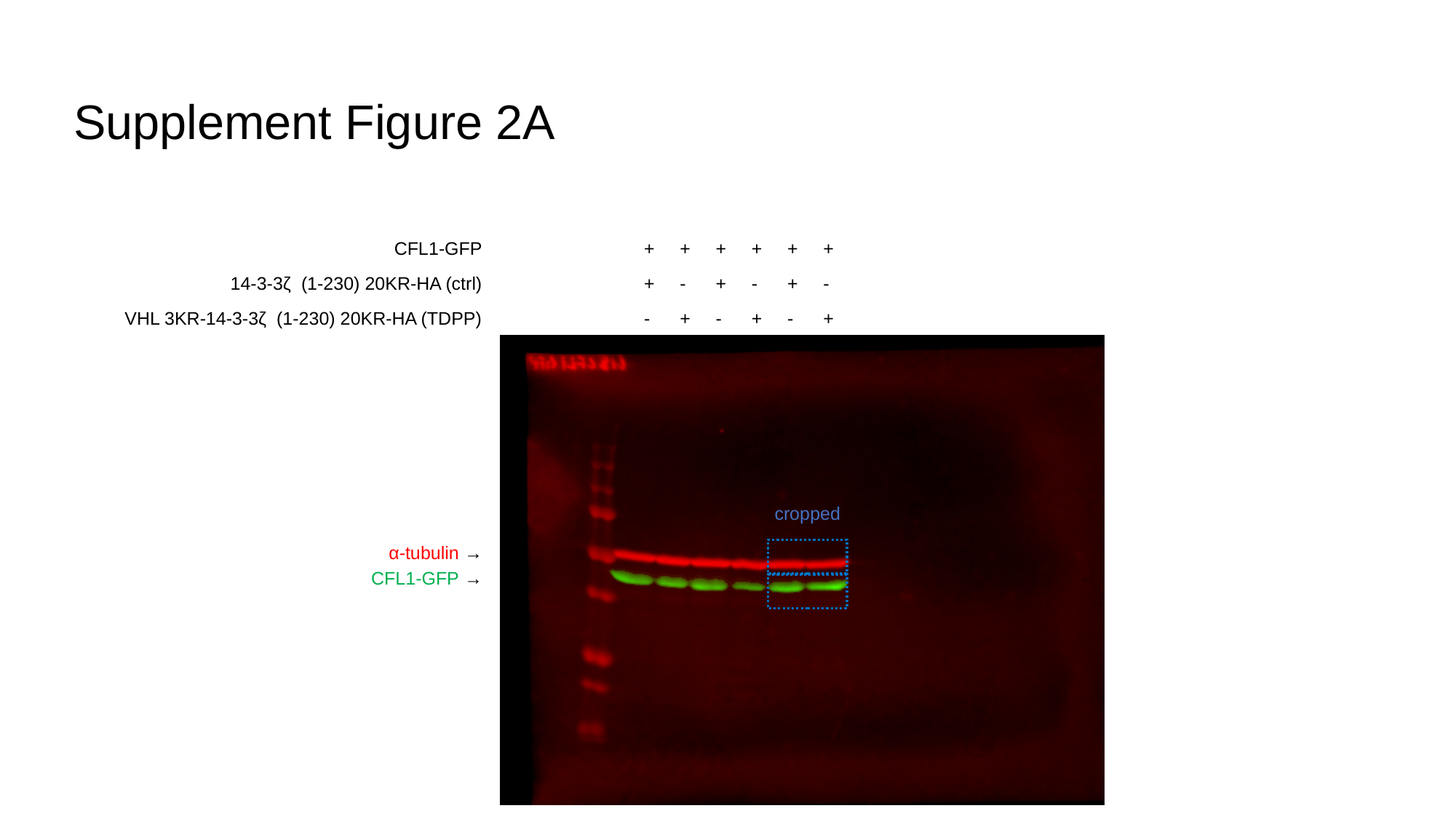

#### Supplement Figure 2A
CFL1-GFP
14-3-3ζ (1-230) 20KR-HA (ctrl)
VHL 3KR-14-3-3ζ (1-230) 20KR-HA (TDPP)
+
+
-
+
-
+
+
+
-
+
-
+
+
+
-
+
-
+
cropped
α-tubulin →
CFL1-GFP →
